## Supplemental data for "Human adipose stromal cells differentiate towards a tendon phenotype with adapted visco-elastic properties in a 3D-culture system"

### Experimental procedures

*Cleaved Caspase 3 immunostaining*

3D-hASCs construct 12µm cryosections were subject to a standard immunofluorescence protocol using human Cleaved Caspase 3 (Cell Signaling; ref 9664S) primary antibody at 1:100 dilution and Goat anti-Rabbit IgG (H+L) Cross-Adsorbed, Alexa Fluor™ 488 (Invitrogen/ Thermo Fisher scientific; ref A11008) secondary antibody at 1:200 dilution. Finally, sections were stained with DAPI (D9542, Sigma-Aldrich) to visualize cell nuclei and phalloidin (A12380 Alexa Fluor Phalloidin, Invitrogen/ ThermoFisher scientific) to visualize cytoskeletal F-actin. Fluorescent images were captured using a Zeiss Axio Observer Z1 microscope equipped with a Zeiss Apotome 2 and an Axiocam 506 monochrome camera.

*Adipogenic Differentiation*

hASCs from each patient were plated at passage 3 at a density of 3,000 cells/cm2 in 6-well plates pre-coated with animal component-free (ACF) cell attachment substrate (Stemcell Technologies; ref 07130). Cells were cultured in ACF Plus medium until confluence. Control cells were cultured in ACF Plus medium (Stemcell Technologies; ref 05446) whereas adipogenic differentiation was tested by replacing the ACF Plus medium with human Mesencult adipogenic differentiation medium (Stemcell Technologies; ref 05412) for 13 days. Medium was changed every 2-3 days.

*Osteogenic Differentiation*

hASCs were plated at passage 3 at a density of 8,000 cells/cm2 in 6-well plates pre-coated with animal component-free (ACF) cell attachment substrate (Stemcell Technologies; ref 07130). Cells were cultured in ACF Plus medium until confluence. Control cells were cultured in ACF Plus medium (Stemcell Technologies; ref 05446) whereas osteogenic differentiation was tested by replacing the ACF Plus medium with human Mesencult osteogenic differentiation medium (Stemcell Technologies; ref 05465) for 14 days. Medium was changed every 4 days.

*Cell Staining*

Adipogenic differentiation was assessed by Oil Red O staining and osteogenic differentiation was assessed by Alizarin Red Staining, as previously described (Waldner et al. 2018; Bléher et al. 2020).

*RNA isolation*

Control and differentiation media were removed and replaced with 350µL RLT in each well or tube. Total RNA was isolated using the RNeasy mini kit (Qiagen; ref 74106), according to the manufacturer's instructions, including a 15 min of DNase I (Qiagen; ref 79254) treatment.

*Reverse-Transcription and quantitative real time PCR*

Total RNAs extracted from control, adipogenic or osteogenic cells were Reverse Transcribed using the High Capacity Retro-transcription kit (Applied Biosystems; ref 4387406). Quantitative PCR analyses were performed using primers listed in Supplementary Table 1 and SYBR Green PCR Master Mix (Applied Biosystems; ref 4385614). The relative mRNA levels were calculated using the $2^{\wedge-\Delta\Delta Ct}$ method (Livak and Schmittgen 2001). The Cts were obtained from Ct normalized to *YWHAZ* level in each sample.


### Figure Legends

**Supplementary Figure 1: The Flexcell bioreactor.** A) This technology consists of a holder with 24 perforated cylindrical moulds connected to a vacuum pump and controlled by computer. B) Four 6-well plates can be placed on these moulds. The bottom of the wells consists of a rubber membrane with two thick anchor points. When the vacuum is applied, the bottom of the wells takes the cylindrical shape of the moulds. hASCs embedded in 3.5% collagen hydrogel are seeded between the two anchor points and incubated at 37°C, 5% $CO_2$ for 2 hours. C) At the end of the 2 hours incubation, the vacuum is broken and the plate is removed from the molds. The plate is left in the incubator at 37°C, 5% $CO_2$ for 48 hours before experimentation (Day -2 timepoint). D) Pictures of 3D-hASC constructs and 3D-no cell constructs not seeded with cells. Scale bars, 0.5 cm. E) Diameters were measured from constructs at Day 0 (n=12), Day 7 (n=12), Day 14 (n=12) and Day 21 (n=12). Each color represents a set of experiments. 2

independent experiments were performed with n=6 biological replicates for each experiment. F) Cross-section areas of 3D-no cell and 3D-hASC constructs were calculated from diameters measurements presented Figure 1B: Day 0 (n=11), Day 7 (n=15), Day 14 (n=14) and Day 21 (n=19). Each color represents a set of experiments. 4 independent experiments were performed with 3<n<8 biological replicates for each experiment. For 3D-hASC constructs, the p-values were obtained using the Mann-Whitney test compared to each following stage. * P<0.05, **** P<0.0001. $^{\#}$ indicates the p-values of cross-section areas of 3D-no cell versus 3D-hASC constructs, $^{\#\#\#\#}$ P <0.0001.

**Supplementary Figure 2. 3D-hASC constructs did not show apoptosis** A) 12µm transverse sections of 3D-hASC constructs on Day 0 and Day 21 of culture were stained with CAS3 antibody to detect apoptotic cells and DAPI/ Phalloidin to visualize cell nuclei and cytoskeletal organization. Scale bars 50 µm. B) Cells migrating from human 3D-hASC constructs. Scale bar: 100µm.

**Supplementary Figure 3. Adipogenic and Osteogenic differentiation of hASCs.** hASCs were subjected to A) Adipogenic differentiation for 13 Days. Control (cultured in basal medium) and differentiated hASCs were stained with Oil Red O and *PPARG* mRNA expression level was analyzed by RT-qPCR. Scale bars 100µm. B) 21-Day 3D-hASC constructs were stained with Oil Red O and *PPARG* mRNA expression level was analyzed by RT-qPCR. Scale bars 20µm. hASCs were subjected to C) Osteogenic differentiation for 14 Days. Control (cultured in basal medium) and differentiated hASCs were stained with Alizarin Red S and *OCN* mRNA expression level was analyzed by RT-qPCR. Scale bars 100µm D) 21-Day 3D-hASC constructs were stained with Alizarin Red S and *OCN* mRNA expression level was analyzed by RT-qPCR. Each differentiation experiment was performed 2 independent times with n ≥ 3 biological replicates. Scale bars 20µm.

**Supplementary Figure 4. *COL3A1* mRNA expression in 3D-hASC constructs.** A) 3D-hASC constructs at Day 7 were transversally cryo-sectioned. 12 µm sections were hybridized with the DIG-labeled antisense probes for *COL3A1*. Scale bar: 500µm. B) 3D-hASC constructs at Day 0, Day 7, Day 14 and Day 21 were transversally cryo-sectioned. 12 µm sections were hybridized with the DIG-labeled antisense probes for *COL3A1*. Scale bars: 50µm.

**Supplementary Figure 5. Cellular and nuclear organization in 3D engineered 3D-hASCs.** A) 3D-hASCs longitudinal section representation. B) Longitudinal sections of 3D-hASC constructs were performed on Day 0, Day 2, Day 4, Day 7, Day 14 and Day 21 3D-constructs and stained with DAPI/Phalloidin to visualize cell shape. Scale bars: 200 µm. White squares represent higher magnification. Scale bars: 100µm.

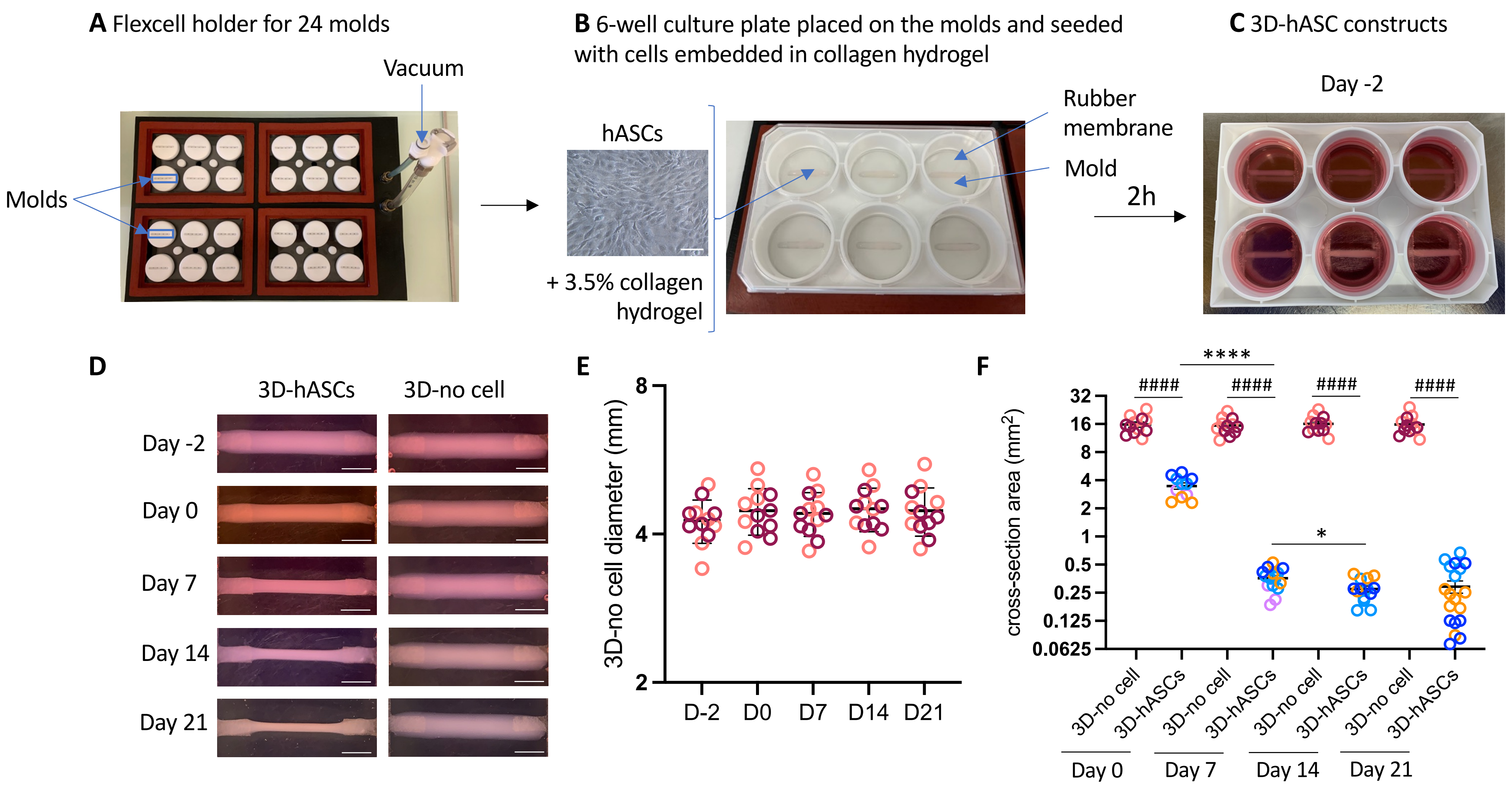


**Supplementary Figure 1**

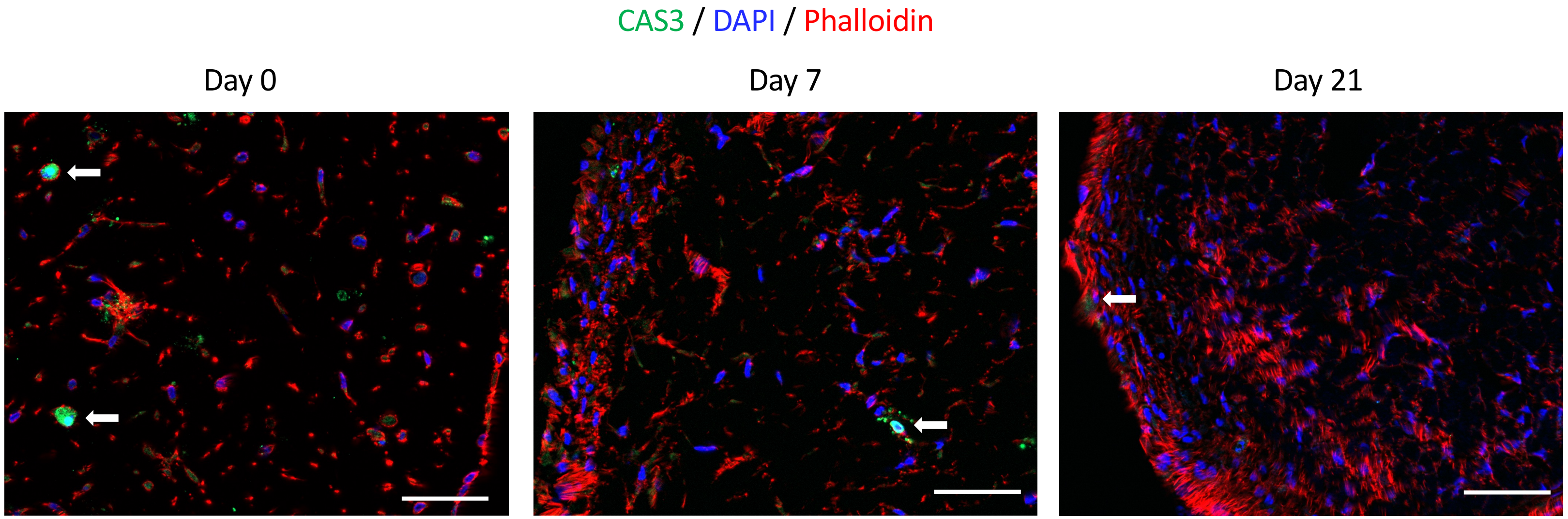


**Supplementary Figure 2**

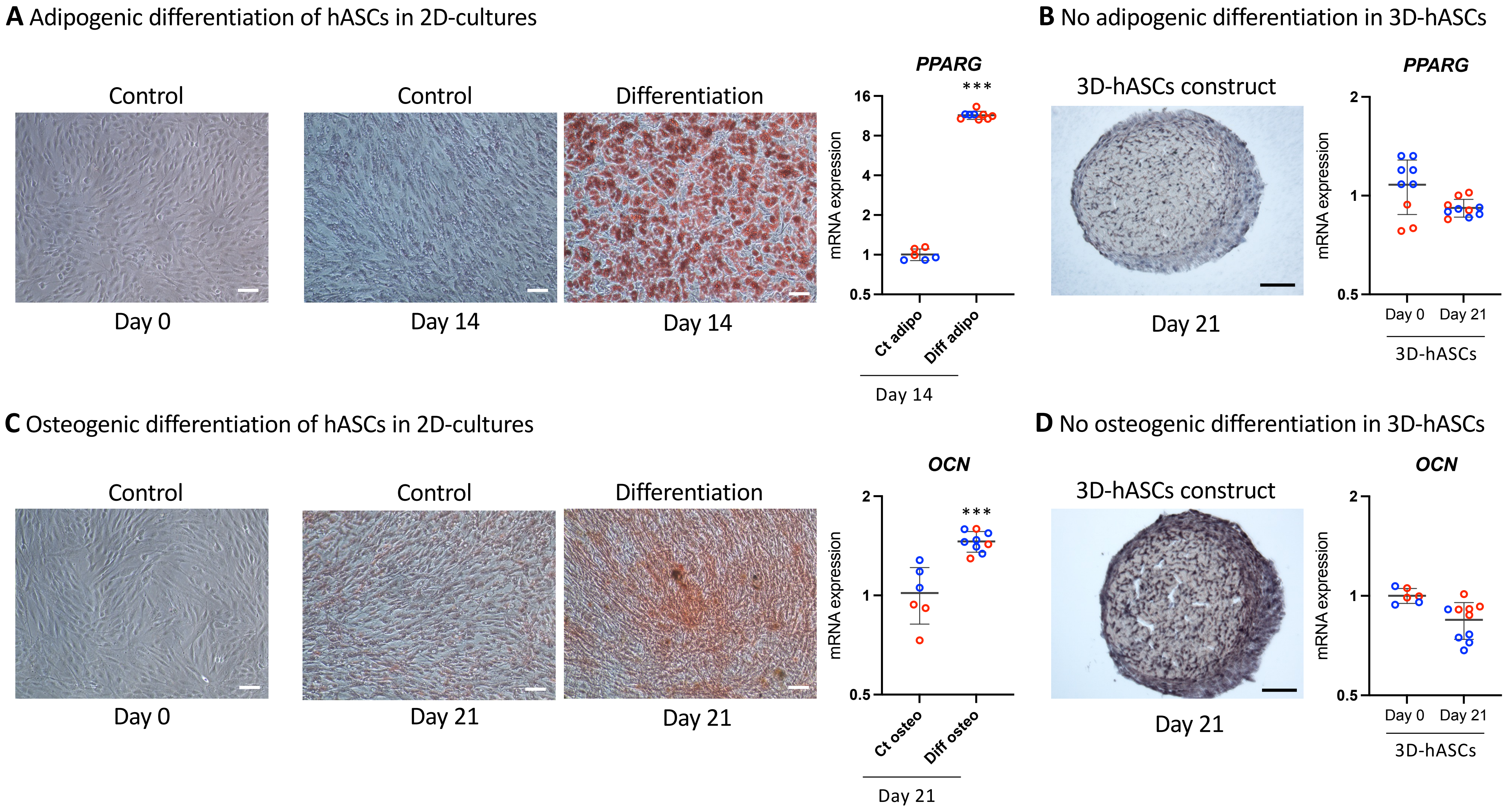


**Supplementary Figure 3**

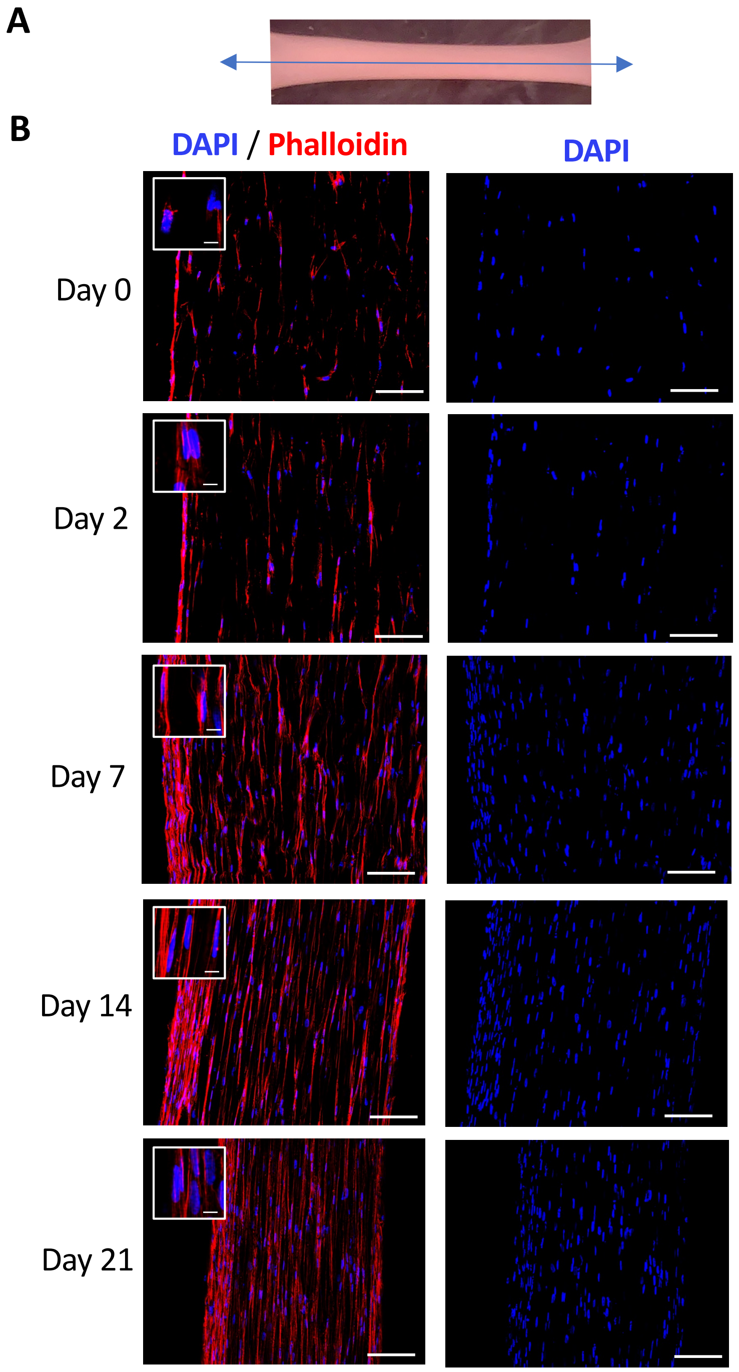


**Supplementary Figure 4**

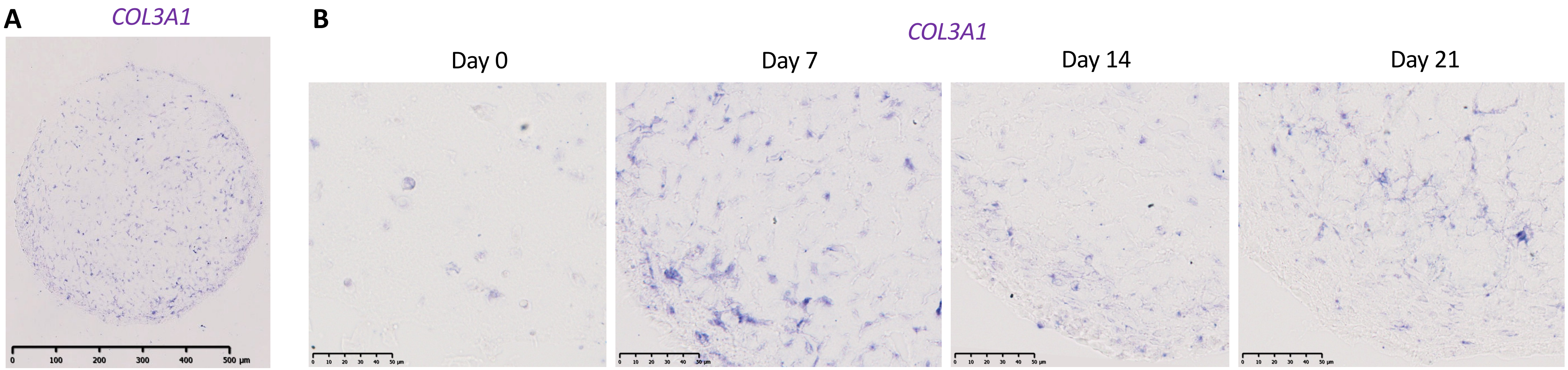


**Supplementary Figure 5**

**Supplementary Table 1**

| | RT-qPCR Forward primer | RT-qPCR Reverse primer | Reference |
|---|---|---|---|
| ***hSCX*** | CCCAAACAGATCTGCACCTT | CGGTCCTTGCTCAACTTTCT | Designed in the lab |
| ***hMKX*** | CATCGTCATCAGAACTGAAGGCA | TCTGTTAGCTGCGCTTTCACCC | (Bayer et al. 2014) |
| ***hTNMD*** | AGCACTTCTGGCCGGAGG | AAGTGTGCTCCATGTCATAGGCT | Designed in the lab |
| ***hTM4SF1*** | CAGCCCTTGGCTTAGCAGAA | ACTCGGACCATGTGGAGGTA | Designed in the lab |
| ***hANXA1*** | AGTTCTTTGCAAGAAGGTAGAGA | CTGATCCGGGACCACCTTTG | Designed in the lab |
| ***hS100A10*** | GGCTACTTAACAAAGGAGGACC | GAGGCCCGCAATTAGGGAAA | Głowacka et al., 2021 |
| ***hCOL1A1*** | GATGGCTGCACGAGTCACAC | GTATTCAATCACTGTCTTGCCCC | Designed in the lab |
| **h*COL6A3*** | AACATCGGCACTTGCCCTTA | ATATCAGCAGCCGCACCATT | Designed in the lab |
| ***hCOL14A1*** | ACTCCGAGGGAAGAGAGCAA | TACATGGGGTGTAGCAGCCA | Designed in the lab |
| ***hDPT*** | TGTCGCTACAGCAAGAGGTG | GTGGTTGTTGCTCCTCGGAT | Designed in the lab |
| ***hCOL2A1*** | TGGCTGACCTGACCTGATGTCC | TGCAGTCTGCCCAGTTCAGGTC | Designed in the lab |
| ***hPOSTN*** | GAGGAAGTTGCAAGCCAACA | CACTGAGAACGACCTTCCCT | Designed in the lab |
| ***hTHBS2*** | GACACGCTGGATCTCACCTAC | GAAGCTGTCTATGAGGTCGCA | (Xu et al. 2020) |

| | | | |
|---|---|---|---|
| ***hPPARG*** | AAGCCCTTCACTACTGTTGACT | CAGGCTCCACTTTGATTG | (Waldner et al. 2018) |
| ***hOCN*** | GGCGCTACCTGTATCAATGG | TCAGCCAACTCGTCACAGTC | Designed in the lab |
| ***hYWHAZ*** | CCGCTGGTGATGACAAGAAAGGGAT | AGGGCCAGACCCAGTCTGATAGGA | (Ragni et al. 2013) |
| Gene name | In situ hybridization probe Forward primer | In situ hybridization probe T7-Reverse primer | Reference |
| ***hSCX*** | GGTCGCTACCTGTACCCTGA | TAATACGACTCACTATAGGGCCTGAGGCAGAAGGTGCAGAT | Designed in the lab |
| ***hTNMD*** | TGGAAATGGCACTGATGAAA | TAATACGACTCACTATAGGGCCAGCATTGGGTCAAATTCAA | Designed in the lab |
| ***hCOL1A1*** | CTCCCCAGCTGTCTTATGGC | TAATACGACTCACTATAGGGCGCACCATCATTTCCACGAGC | Designed in the lab |
| ***hCOL3A1*** | CCTACTCGCCCTCCTAATGG | TAATACGACTCACTATAGGGCTCGAAGCCTCTGTGTCCTTT | Designed in the lab |
| ***hTHBS2*** | ACCAGGACAAAGACACGACC | TAATACGACTCACTATAGGGCCCACGTACATCCGGCTCTTT | Designed in the lab |

**Supplementary Table 1 references**


Bayer, Monika L., Peter Schjerling, Andreas Herchenhan, Cedric Zeltz, Katja M. Heinemeier, Lise Christensen, Michael Krogsgaard, Donald Gullberg, and Michael Kjaer. 2014. "Release of Tensile Strain on Engineered Human Tendon Tissue Disturbs Cell Adhesions, Changes Matrix Architecture, and Induces an Inflammatory Phenotype." *PLoS ONE* 9 (1). https://doi.org/10.1371/journal.pone.0086078.

Ragni, Enrico, Mariele Viganò, Paolo Rebulla, Rosaria Giordano, and Lorenza Lazzari. 2013. "What Is beyond a QRT-PCR Study on Mesenchymal Stem Cell Differentiation Properties: How to Choose the Most Reliable Housekeeping Genes." *Journal of Cellular and Molecular Medicine* 17 (1): 168–80. https://doi.org/10.1111/j.1582-4934.2012.01660.x.

Waldner, Matthias, Wensheng Zhang, Isaac B. James, Kassandra Allbright, Emmanuelle Havis, Jacqueline M. Bliley, Aurora Almadori, et al. 2018. "Characteristics and Immunomodulating Functions of Adipose-Derived and Bone Marrow-Derived Mesenchymal Stem Cells across Defined Human Leukocyte Antigen Barriers." *Frontiers in Immunology* 9 (JUL). https://doi.org/10.3389/fimmu.2018.01642.

Xu, Chunjie, Lei Gu, Manzila Kuerbanjiang, Siyuan Wen, Qing Xu, and Hanbing Xue. 2020. "Thrombospondin 2/Toll-Like Receptor 4 Axis Contributes to HIF-1α-Derived Glycolysis in Colorectal Cancer." *Frontiers in Oncology* 10 (November). https://doi.org/10.3389/fonc.2020.557730.
